# Mitigating sTNF/TNFR1 activation on VGluT2+ spinal cord interneurons improves immune function after mid-thoracic spinal cord injury

**DOI:** 10.1101/2024.07.09.602690

**Authors:** Tetyana Martynyuk, Jerome Ricard, Valerie Bracchi-Ricard, Samuel Price, Jenna McGrath, Kimberly Dougherty, Veronica Tom, John R. Bethea

## Abstract

Spinal cord injury (SCI) is a devastating condition with 250,000 to 500,000 new cases globally each year. Respiratory infections, e.g., pneumonia and influenza are the leading cause of death after SCI. Unfortunately, there is a poor understanding of how altered neuro-immune communication impacts an individual’s outcome to infection. In humans and rodents, SCI leads to maladaptive changes in the spinal-sympathetic reflex (SSR) circuit which is crucial to sympathetic function. The cause of the impaired immune function may be related to harmful neuroinflammation which is detrimental to homeostatic neuronal function, aberrant plasticity, and hyperexcitable circuits. Soluble tumor necrosis factor (sTNF) is a pro-inflammatory cytokine that is elevated in the CNS after SCI and remains elevated for several months after injury. By pharmacologically attenuating sTNF in the CNS after SCI we were able to demonstrate improved immune function. Furthermore, when we investigated the specific cellular population which may be involved in altered neuro-immune communication we reported that excessive TNFR1 activity on excitatory INs promotes immune dysfunction. Furthermore, this observation is NF-kB dependent in VGluT2+ INs. Our data is the first report of a target within the CNS, TNFR1, that contributes to SCI-induced immune dysfunction after T9-SCI and is a potential avenue for future therapeutics.

## Introduction

Spinal cord injuries are devastating, leaving individuals paralyzed below the level of injury. However, the motor impairments faced are not the only hurdles for spinal cord injury patients. Every organ in the body is innervated through the spinal cord via the sympathetic system, thus, patients are faced with a myriad of complications including increased risk for cardiovascular diseases, metabolic syndrome, autonomic dysreflexia, and decreased immune function (DiSabato et al., 2024). In fact, unlike the general population of which 3% succumb to infections, more than 25% of deaths following SCI are due to infections such as influenza, pneumonia, and septicemia. (Center, 2023; Kopp et al., 2023; Soden et al., 2000) This comes from a neurogenic immune deficiency and not merely from a post-traumatic immune deficiency as patients with vertebral fractures not affecting their spinal cord do not exhibit increased susceptibility to infection (Kopp et al., 2023). Pre-clinical studies have established that the disconnection of the sympathetic system from supraspinal control leads to profound changes in the spleen and dramatically affects its function. However, clinical and pre-clinical studies have shown that even lower thoracic SCI result in increased susceptibility to infections (Bracchi-Ricard et al., 2016; Held, Steward, Blanc, & Lane, 2010; Kopp et al., 2023)

High thoracic injuries not only affect the immune system but also the innervation of the lungs, thereby contributing to the enhanced susceptibility to infections as patients cannot breathe or cough properly. After lower thoracic SCI (i.e., T9-SCI), innervation of the lungs is preserved, yet increased susceptibility to infection is well documented. It is also admitted that the supraspinal control over the sympathetic preganglionic neurons (SPNs) that regulate the sympathetic outflow to the spleen is mostly preserved with lower SCI. However, we and others have shown deficiency in anti-viral immunity and increased mortality in mice with lower thoracic T9 SCI (Bracchi-Ricard et al., 2016; Held et al., 2010). It’s crucial to recognize and understand the increased risk for infection that this patient population faces.

Although the descending control over the SPNs is mostly preserved, a pro-inflammatory environment in the spinal cord around SPNs could affect their function and the spinal interneurons synapsing onto them. Spinal cord interneurons (INs) are a diverse group of neurons (Zavvarian, Hong, & Fehlings, 2020) that play critical roles in modulating spinal circuits. Extensively studied in the context of locomotion, INs participate in the regulation of sensory and autonomic circuitry (Zholudeva et al., 2021). Following SCI, IN activation contributes to neuroplasticity (Zholudeva et al., 2021). Increased numbers of synaptic contacts between excitatory VGluT2+ INs and SPNs following high T3 thoracic SCI renders the spinal-splenic circuit more sensitive to activation leading to immune suppression (Ueno, Ueno-Nakamura, Niehaus, Popovich, & Yoshida, 2016). Silencing VGluT2+ INs, using DREADD technology, improved immune competency in T3 SCI mice (Ueno et al., 2016). Of note, in this study the authors did not observe an increase in the number of synapses between VGluT2+ IN and SPNs with a lower thoracic injury but it is unknown if the pro-inflammatory environment due to the injury would affect the excitatory/inhibitory balance in the spinal cord and affect the excitability of INs.

Soluble tumor necrosis factor (sTNF) is one of the first pro-inflammatory cytokines expressed following SCI and is persistently elevated for more than 3 weeks in the spinal cord (Hellenbrand et al., 2021). sTNF, in the CNS, is also known as a factor modulating synaptic plasticity (Heir & Stellwagen, 2020) Elevated levels of sTNF have been correlated with various types of diseases such as cancer, HIV and AIDS, systemic lupus erythematosus, multiple sclerosis, and liver diseases (Diez-Ruiz et al., 1995). In addition, inflammatory environments in the spinal cord persistently activate interneurons, possibly leading to maladaptive sympathetic alterations in the modulation of peripheral organs (Jin et al., 2023; Massey et al., 2024; Ueno et al., 2016; Zavvarian et al., 2020; Zholudeva et al., 2021). However, how chronic inflammation after T9-SCI negatively impacts the immune system is largely unknown. We hypothesized that the pro-inflammatory microenvironment arising within the spinal cord following T9 SCI could alter the function of interneurons and thus disrupt the sympathetic output tone to the spleen.

Specifically, that sTNF signaling through TNFR1 and NF-κB in excitatory, VGluT2+, interneurons could lead to an increase in excitatory input to the sympathetic preganglionic neurons, resulting in worsened antiviral immunity due to dysfunction of the spleen.

To address this hypothesis, we utilized a well-established moderate T9 contusion model (Bracchi-Ricard et al., 2016). First, we pharmacologically inhibited TNFR1 signaling within the injured spinal cord using continuous administration of XPro1595, a selective inhibitor of sTNF, and restored antiviral immunity of mice challenged with live influenza after chronic T9-SCI. Next, we showed that modulating TNFR1 signaling specifically in VGlut2+ excitatory or VGat+ inhibitory neurons reveal an interesting dichotomy of the effects on anti-viral immunity. Furthermore, we sought to evaluate if these effects were NF-kB dependent by inhibiting the activation of NF-kB specifically in VGluT2+ and VGat+ INs and reported that our improvement in anti-viral immunity is NF-kB dependent in VGluT2+ INs. Taken together, our data show that chronic inflammation in the spinal cord after T9-SCI may lead to immune dysfunction by altering the activity of VGluT2+ and VGat+ INs in the sympathetic innervation of the spleen. Our results lead to a potential novel therapeutic strategy for SCI-induced immune dysfunction.

## Results

### Intrathecal inhibition of sTNF restores anti-viral immune function in spinal cord injured mice

Chronic inflammation after SCI has documented roles in various secondary conditions including immune dysfunction (Boehl et al., 2022; Mokhtari & Uludag, 2024; Ryan et al., 2024; Zhang et al., 2024). We have previously shown that chronic T9 SCI negatively impacts anti-viral immunity (Bracchi-Ricard, 2016) (Norden, Qatanani, Bethea, & Jiang, 2019) but how sTNF may contribute to immune dysfunction after T9-SCI remains largely unknown. To investigate the impact of elevated intraspinal sTNF on antiviral immunity following chronic T9 SCI we delivered XPro1595, an anti-sTNF biologic, continuously for 4 weeks via an osmotic pump, directly to the site of injury immediately after SCI (Fig. 1a). Four weeks later, mice were infected intranasally with live influenza particles and monitored daily for body weight loss (Fig. 1b). SCI mice in the saline-treated group lost significantly more weight (a key sign of sickness) than uninjured control mice (Fig. 1b). However, SCI mice treated with XPro1595 lost less weight and weight loss was comparable to that of the uninjured group suggesting improved immune function (Fig. 1b). We determined the amount of residual viral load in the lungs 9-days post-infection by quantitative RT-PCR for the M1 gene encoding a viral matrix protein of influenza. Mice treated with XPro1595 had significantly less residual viral load compared to vehicle-treated SCI mice further suggesting that inhibiting TNFR1 activation in the spinal cord is beneficial for mounting an appropriate immune response after chronic T9 SCI (Fig. 1c). Whereas a thoracic injury at vertebral level T9 does not directly impact the innervation of the lungs it does impact the innervation of the spleen. Upon investigation of the spleen after SCI, we noted a significant decrease in spleen weight and total splenocyte count, excluding RBCs. With the administration of XPro1595, there was a significant increase in both spleen weight and total splenocyte numbers compared to vehicle controls (Fig. 1d, e). Cellular immunity response is well characterized following influenza infection in an uninjured individual, however, the impact of excessive sTNF on the impaired CD8+ T cell response after T9-SCI is largely unknown. CD4+ and CD8+ T cells are two classes of lymphocytes which are crucial for viral clearance and prolonged immunity from subsequent infection (Hufford, Kim, Sun, & Braciale, 2015). We have previously demonstrated that T9-SCI leads to impaired CD8 T cell response to influenza and have increased markers for functional exhaustion (Bracchi-Ricard et al., 2016; Zha, Smith, Andreansky, Bracchi-Ricard, & Bethea, 2014) . However, how increased sTNF in the spinal cord after injury contributes to the mechanism of decreased immune response in the spleen is unknown. Mice which received XPro1595 after SCI had an increased number of CD4+ T cells, CD8+ T cells, as well as activated T Cells (CD44+CD62L^low^) in the spleen (Fig. 1f). XPro1595 infusion in the spinal cord significantly increases virus specific CD8+ T cells in the lungs (Fig. 1g), which have been shown to directly impact the ability of an organism to clear the IAV infection(Cheung et al., 2002; de Jong et al., 2006; Sun, Madan, Karp, & Braciale, 2009). Our results suggest that inhibiting chronic inflammation in the spinal cord modulates the cellular profile of a peripheral organ, the spleen.

**Figure 1.**
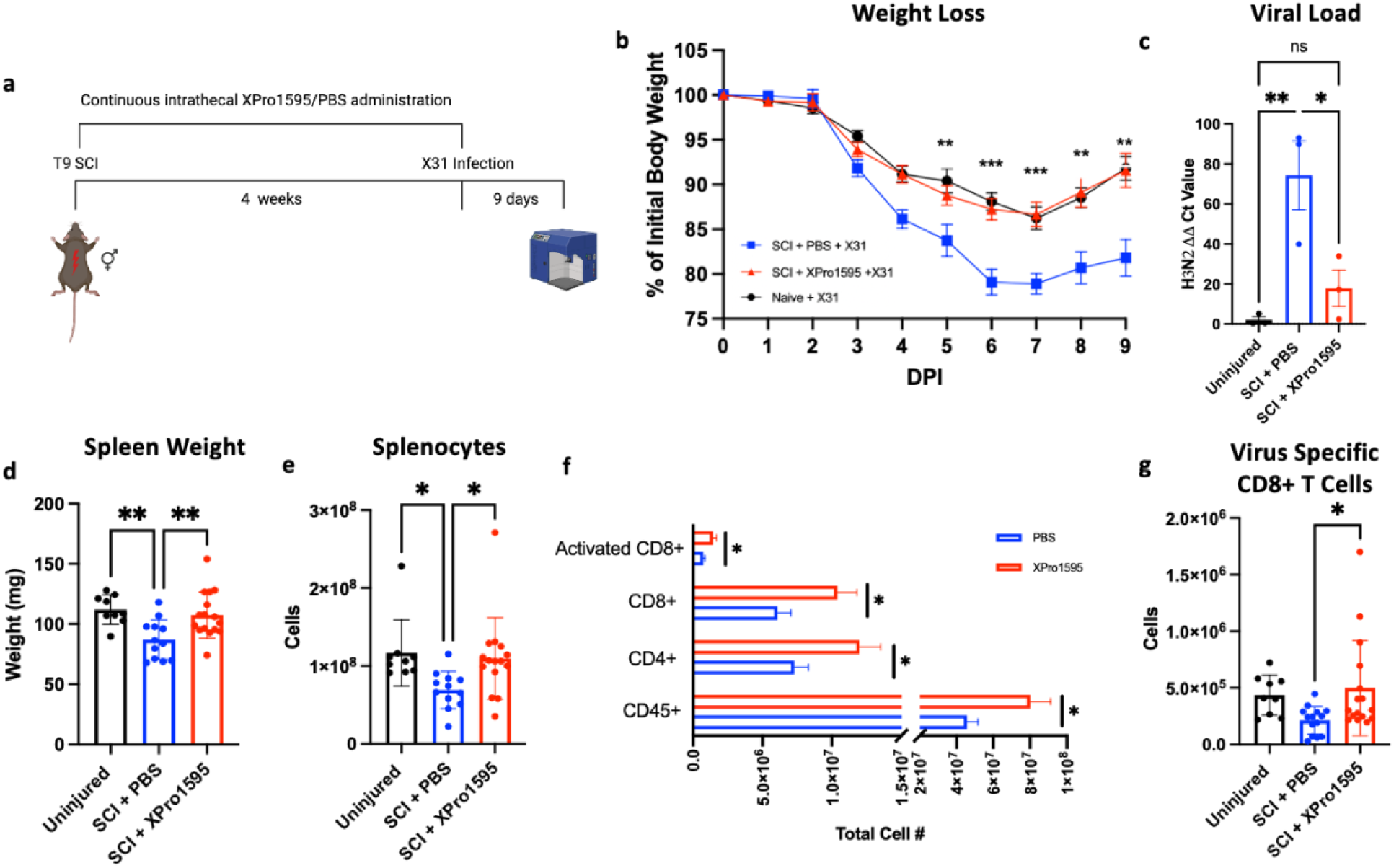
Intrathecal administration of XPro1595 mitigates T9 SCI-induced immune dysfunction. **a** Schematic of timeline used for intrathecal administration and chronic SCI (created with biorender.com) **b** Inhibiting sTNF activity in the spinal cord decreases body weight loss after infection (Naïve n=9, SCI PBS treated n=12, SCI XPro1595 treated n=16, p values represent significance between XPro1595 and PBS treated SCI groups, **** p<0.0001, *** p=0.0002, 8, 9 DPI p=0.0001 using one-way ANOVA, mean ± SEM represented). **c** Quantitative real-time PCR of viral load in lung at 9 DPI with significant increase in ability to clear viral particles with XPro1595 treatment (Uninjured n=4, PBS n=3, XPro1595 n=4, uninjured to PBS ** p=0.0053, PBS to XPro1595 ** p=0.097 using ANOVA, mean ± SEM represented) **d** Significant decrease in spleen atrophy after T9 SCI XPro1595 treatment (Uninjured n=9, PBS n=12, XPro1595 n=16, uninjured to PBS ** p=0.0053, PBS to XPro1595 ** p=0.0084 using ANOVA, mean± SEM represented) **e** Total splenocytes counted at the time of sample collection (Uninjured to PBS * p=0.0407, PBS to XPro1595 * p=0.0496 ANOVA, mean± SEM represented) **f** XPro1595 administration significantly increased CD45+, CD4+, CD8+ and activated (CD44+CD62L+) CD8+ T cells in the spleen (CD45 * p=0.0192, CD4 * p=0.0239, CD8 * p=0.0216, Activated * p=0.0186, using students T Test, mean± SEM represented) and **g** virus specific CD8+ T cells at the primary site of infection, the lung (PBS to XPro1595 * p=0.0307 ANOVA mean± SEM represented).

### TNFR1 KD on excitatory (VGluT2+) INs improves anti-viral immunity after T9 SCI

Previous work from our lab and others has highlighted the role played by spinal interneurons in modulating sympathetic circuitry output. We showed that high thoracic (T3) SCI specifically increases the activity of thoracic interneurons in the sympathetic systems compared to sympathetic preganglionic neurons (Mironets et al., 2020; Ueno et al., 2016) (Mironets et al., 2018). To test if the persistent elevation of sTNF in the spinal cord after SCI negatively impacts immune function via effects on excitatory interneurons, we knocked down TNFR1 specifically on VGluT2+ INs. To investigate if absence of TNFR1 on excitatory interneurons results in an improved anti-viral immunity after T9 SCI we used homozygous TNFR1 f/f mice which were injected intraspinally with AAV5 expressing Cre under the VGluT2 promoter (from here on annotated as TNFR1^VGlut2+^) (Fig. 2a). We used RNAscope technology to verify that our experimental approach specifically down-regulated TNFR1 in VGluT2+ INs. Specifically, we analyzed the number of cells positive for both TNFR1 and VGluT2+ before and after genetic manipulation (Fig. 2b). We noted a third reduction in TNFR1 expression compared to EV controls (Fig. 2c). For these experiments, mice were administered AAV5 injections 3 weeks prior to T9 SCI. Four weeks after T9-SCI mice were infected intranasally with live influenza articles and euthanized for cellular analysis 9 days after infection (Fig. 2a). When challenged with live virus after chronic T9-SCI, TNFR1^VGlut2^ mice lost less weight than the EV controls (Fig. 2d). In addition, TNFR1^VGlut2+ KD^ mice had significantly less viral load remaining in their lung than TNFR1^EV^ controls and a comparable amount to uninjured infected mice indicating that excessive TNFR1 activation on VGluT2^+^ interneurons after SCI by sTNF negatively impacts anti-viral immunity (Fig. 2e). TNFR1^VGlut2^ also mitigated splenic atrophy (Fig. 2f), however it did not impact the total amount of splenocytes (Fig. 2g). Although we report this significant impact on the reduction of body weight after influenza infection, we did not notice any differences in CD45+, CD4+, CD8+, or activated T cells in the spleen (Fig. 2h). Furthermore, virus-specific CD8+ T cells in the lung were not increased with KD of TNFR1 on VGluT2+ INs (Fig. 2i). Our results indicate that TNFR1+ activation on INs after T9-SCI improves anti-viral immunity independent of the abundance of T cells in the spleen.

**Figure 2.**
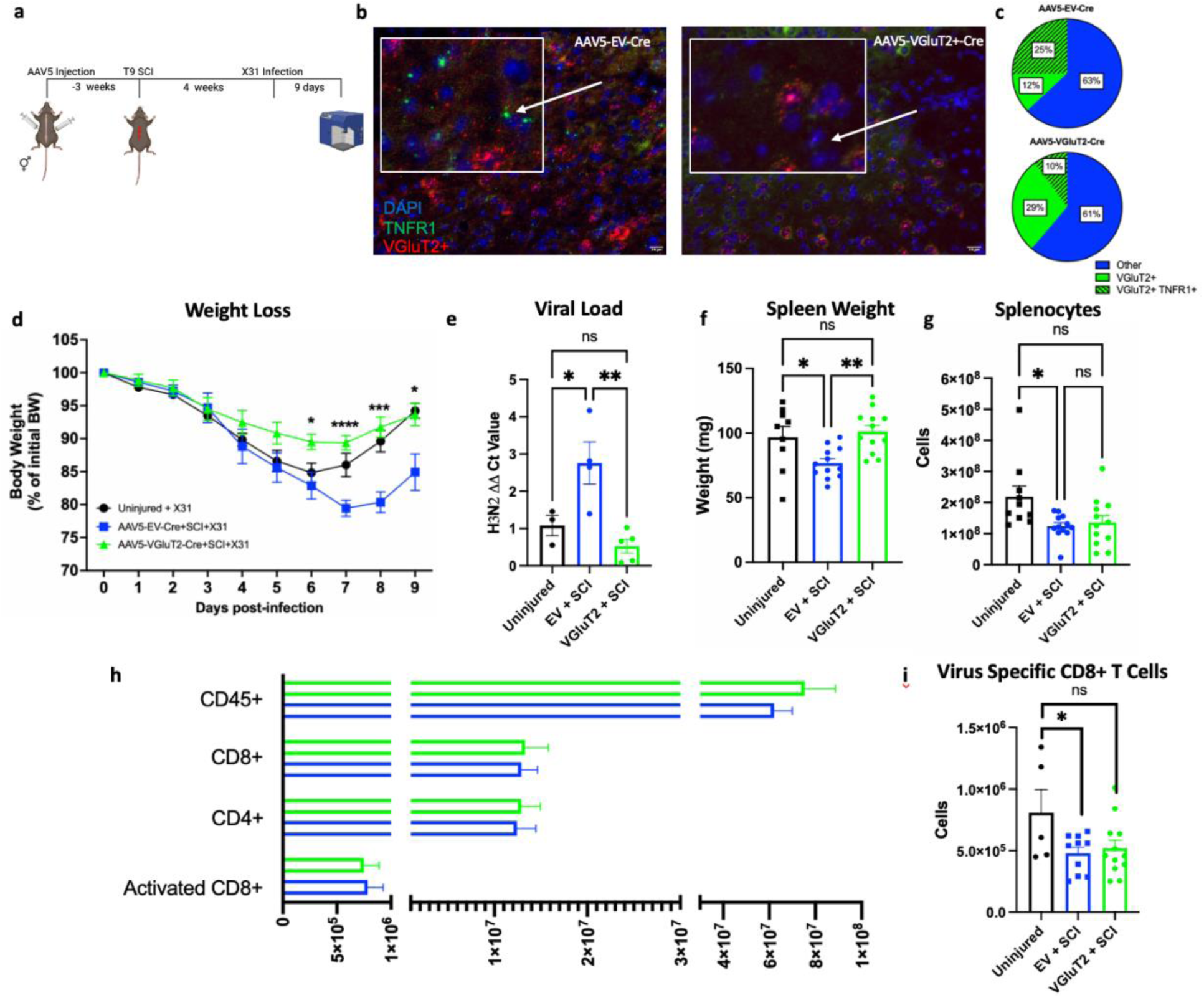
TNFR1 KD on VGluT2+ INs in the thoracic SC improves anti-viral immunity after chronic T9-SCI. **a** Timeline showing AAV5-VGluT2-Cre or EV-Cre administration 3 weeks before T9 SCI. Injections indicate the microinjections made to each side of the thoracic spinal cord for site-specific Cre delivery Chronic T9-SCI was followed by intranasal infection of live influenza virus (H3N2 X31). **b, c** Representative images of RNAscope analysis of TNFR1+ cells in the thoracic SC with EV or VGluT2-Cre administration and quantification of successful KD. DAPI cells were counted, followed by VGluT2+ and TNFR1+. Chart represents the proportion of VGluT2+ cells (green) VGluT2+TNFR1+ (dashed green) and all other cells (blue). **d** Body weight loss of mice for 9 DPI after chronic T9-SCI. (Uninjured + X31 n=10 AAV5-EV-Cre n=8, AAV5-VGluT2-Cre n=12, p values represent significance between AAV5-EV and AAV5-VGluT2, 6 DPI * p= 0.0366, 7 DPI **** p=<0.0001, 8 DPI *** p=0.002, 9 DPI * p=0.0483 using ANOVA, mean ± SEM represented) **e** Viral load remaining in the lung at 9 DPI as determined by RT-qPCR for H3N2 viral load (Uninjured + X31 n=3, AAV-EV-Cre n=4, AAV5-VGluT2-Cre n=5, uninjured to AAV5-EV-Cre *p=0.0392, AAV5-EV to VGluT2-Cre ** p=0.0039 using ANOVA, mean ± SEM represented) **f** Spleen weight of mice after 9 DPI weighed at the time of sample collection and processing for flow cytometry analysis. (Uninjured to AAV5-EV-Cre * p=0.0454, AAV5-EV to VGluT2-Cre ** p=0.0052 ANOVA, mean± SEM represented) **g** Total splenocytes counted at the time of sample collection (Uninjured to PBS * p=0.0407, PBS to XPro1595 * p=0.0496 ANOVA, mean± SEM represented) **h** Bar graph representing the mean± SEM of CD45+, CD4+, CD8+ and CD44+CD62Llow leukocytes, all cells were gated from single, live, CD45+. **i** virus specific CD8+ T cells at the primary site of infection, the lung (AAV-EV to VGluT2-Cre ns p=0.6392, using student’s t test, mean± SEM represented).

### TNFR1 KD on inhibitory (VGat+) INs does not impact anti-viral immunity after T9 SCI

Inhibitory, VGat+, interneurons have critical functions in mediating hyperexcitability in the brain (Clatot et al., 2024; Lee & Maguire, 2013), and silencing VGat+ causes detrimental effects during development and disease. Thus, we hypothesized that proper TNFR1 function on VGat+ interneurons in the thoracic spinal cord were necessary to maintain proper immune function. Using the previously mentioned timeline (Fig. 2a) we knocked down TNFR1 on VGat+ INs using an AAV5-VGat-Cre, here on referred to as TNFR1^VGat^. AAV5-VGat-Cre administration yielded a 45% knockdown of TNFR1 on VGat+ INs (Fig. 3a, b). TNFR1^VGat+^ did not have improved anti-viral immunity after chronic T9-SCI and mice lost the same amount of body weight as the empty vector controls, TNFR1^EV^ (Fig. 3b). TNFR1^VGat^ had significantly more viral load remaining in their lungs 9 DPI (Fig. 3d). In addition, TNFR1^VGat^ did not have any beneficial effects on the spleen weight, total splenocytes, or immune cells in the spleen indicating that decreasing the activity of inhibitory interneurons in the sympathetic circuit may be harmful rather than beneficial to secondary conditions such as immune dysfunction (Fig. 3 e-g).

**Figure 3.**
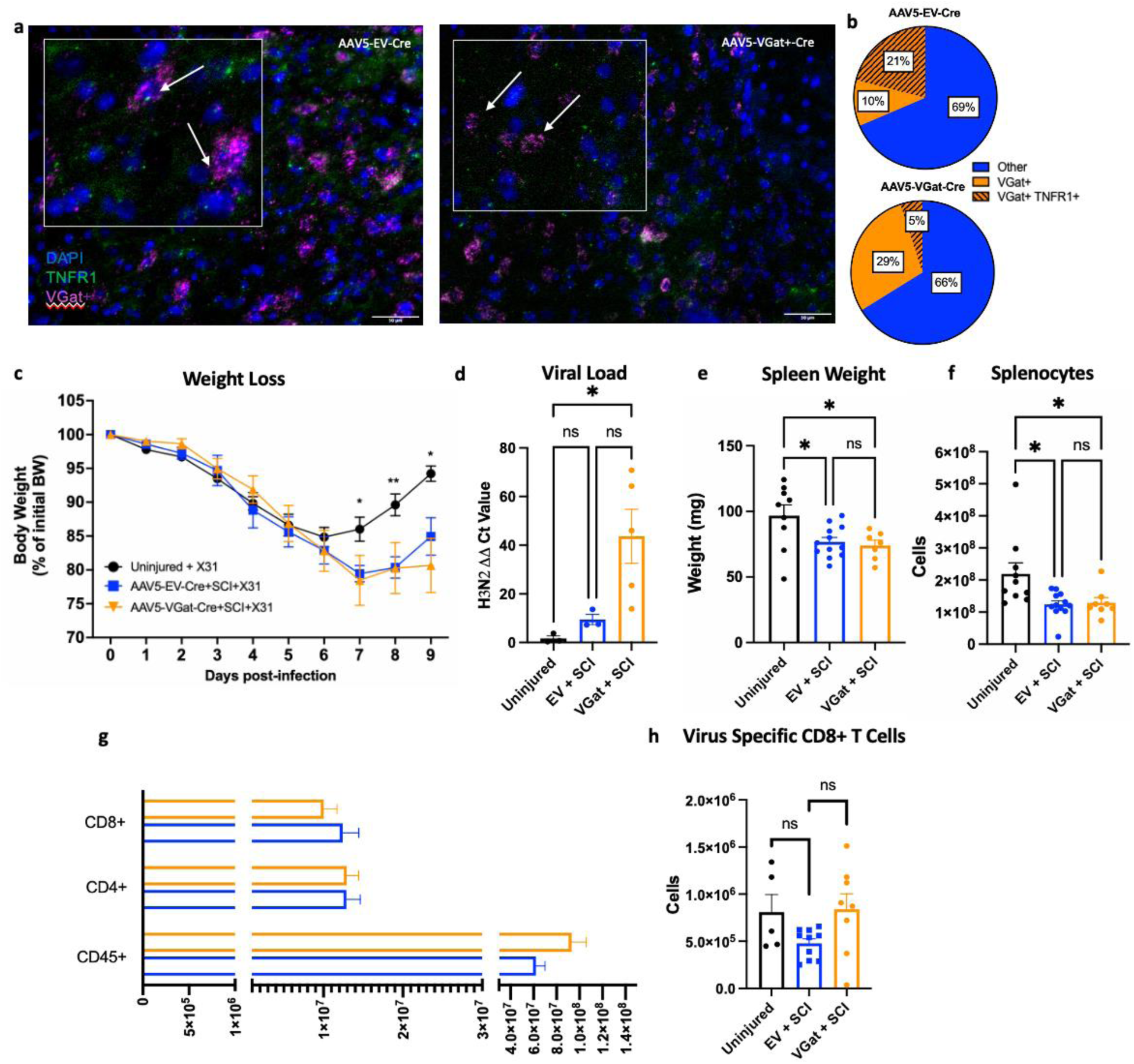
TNFR1 KD on VGat+ INs does not improve anti-viral immunity after chronic SCI despite an increase in virus-specific CD8+ T cells in the lung. **a, b** Representative images of RNAscope analysis of VGat+ TNFR1+ cells in the thoracic SC with EV or VGat-Cre administration and quantification of successful KD. DAPI cells were counted, followed by VGat+ and TNFR1+. Charts are made representing (orange) total VGat+ out of total (blue) and (dashes orange) VGat+TNFR1+. **c** Body weight loss of mice for 9 DPI after chronic T9-SCI (Uninjured + X31 n=10 AAV5-EV-Cre n=8, AAV5-VGat-Cre n=8, p values represent significance between AAV5-EV and AAV5-VGat, 7 DPI *p= 0.0236, 8 DPI ** p=0.0027, 9 DPI * p=0.0356, using ANOVA, mean ± SEM represented). **d** Viral load remaining in the lung at 9 DPI as determined by RT-qPCR for H3N2 viral load (Uninjured + X31 n=3, AAV-EV-Cre n=4, AAV5-VGat-Cre n=5, uninjured to VGat-Cre *p=0.0280 using ANOVA, mean ± SEM represented). **e, f** Spleen weight and total live splenocyte count of spleens from mice after 9 DPI. Weighed and analyzed at the time of sample collection and processing for flow cytometry analysis (Spleen weight, uninjured to AAV5-EV-Cre * p=0.0454, AAV5-EV to VGluT2-Cre ** p=0.0467 ANOVA, mean± SEM represented. Splenocyte number, uninjured to AAV5-EV-Cre * p=0.0133, AAV5-EV to VGat-Cre *p=0.384 using ANOVA, mean ± SEM represented). **g** Bar graph representing the differences of CD45+, CD4+, and CD8+ leukocytes All cells were gated from single, live, CD45+. **h** virus specific CD8+ T cells at the primary site of infection, the lung (AAV-EV to VGat-Cre * p=0.0350, using student’s t test, mean± SEM represented).

### Beneficial effects of TNFR1 KD on VGluT2+ INs is Nf-kB dependent

NF-kB is the canonical downstream pathway of TNFR1, and its transcriptional targets promote inflammation and excitability in various cell types including immune cells and neurons (Liu, Zhang, Joo, & Sun, 2017; Yu, Lin, Zhang, Zhang, & Hu, 2020). Due to our findings that TNFR1 activation on VGluT2+ INs negatively impacts anti-viral immunity, we sought to investigate how NF-κB activation in VGluT2+ and VGat+ INs contributes to T9-SCI-induced immune dysfunction. Using IKKβ ^f/f^ mice we utilized the same AAV5-VGluT2-Cre and AAV5-VGat-Cre to block NF-kB activation by inhibiting the ability of IKK to phosphorylate IκBa, hence preventing its degradation, leading to the sequestration of NF-κB dimers in the cytoplasm (Fig. 4a). We hypothesized that persistent NF-κB activation through TNFR1 would lead to persistent activation in the sympathetic circuit resulting in a negative impact on the spleen, increasing inflammation and dampening anti-viral immunity. Successful knock-down (Fig. 4b,c) of IKKβ in VGluT2+INs, here on referred to as IKKβ ^VGluT2^ improved viral clearance after chronic T9-SCI when challenged with live influenza although mice did not lose significantly less weight until 9 days post infection (Fig. 4d, e). Unlike TNFR1^VGluT2^, IKKβ^VGluT2^ did not impact spleen weight (Fig. 4f). However, splenocytes and CD45+ splenocytes were significantly lower than uninjured controls suggesting that IKKβ ^VGluT2^ may impact function of splenocytes rather than their abundance (Fig. g, h).

**Figure 4.**
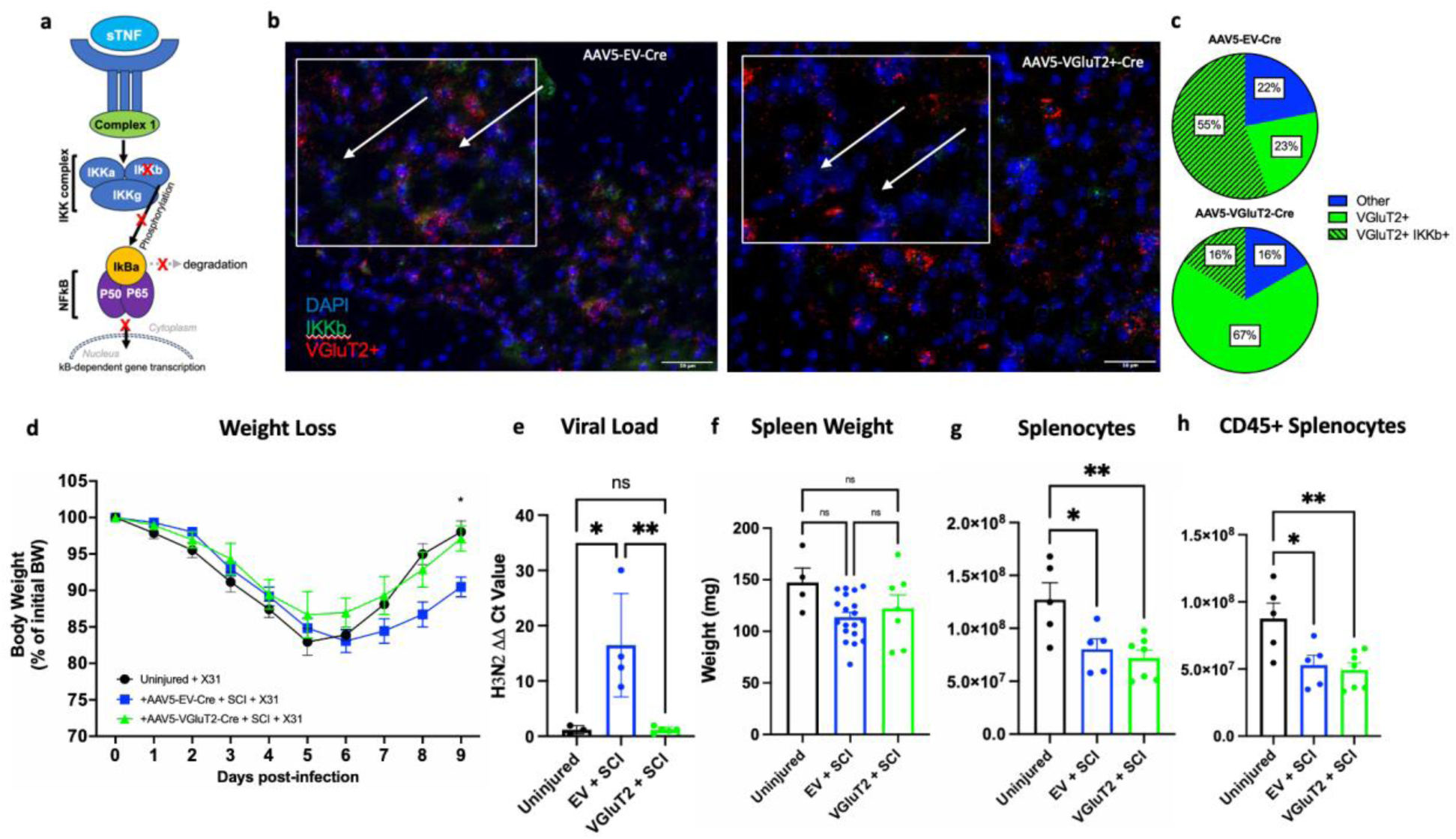
IKKβ KD on VGluT2+ INs improves anti-viral immunity after chronic SCI independent of the abundance of lymphocytes in the spleen. **a** Cartoon representative of the manipulation of IKKβ knock-down **b, c** Representative images of RNAscope analysis of VGluT2+ IKKβ cells in the thoracic SC with EV or VGluT2+-Cre administration and quantification of successful KD. Charts are made representing (green) total VGluT2+ out of total (blue) and (dashed green) VGluT2+ IKKβ +. **d** Body weight loss of mice for 9 DPI after chronic T9-SCI (Uninjured + X31 n=7 AAV5-EV-Cre n=20, AAV5-VGluT2-Cre n=20, p values represent significance between AAV5-EV and AAV5-VGat, 9 DPI * p=0.0288, using ANOVA, mean ± SEM represented). **e** Viral load remaining in the lung at 9 DPI as determined by RT-qPCR for H3N2 viral load (Uninjured + X31 n=3, AAV-EV-Cre n=4, AAV5-VGluT2-Cre n=5, uninjured to EV-Cre * p=0.0124, AAV5-EV to VGluT2-Cre ** p=0.0089 using ANOVA, mean ± SEM represented). **f,g** Spleen weight and total live splenocyte count of spleens from mice after 9 DPI. Weighed and analyzed at the time of sample collection and processing for flow cytometry analysis (total splenocyte uninjured + X31 to AAV-EV * p=0.0282, uninjured + X31 to VGluT2-Cre ** p=0.0043, using ANOVE, mean ± SEM represented). **h** Bar graph representing the total amount of CD45+ splenocytes in the spleen (uninjured + X31 to AAV-EV * p=0.0211, uninjured + X31 to VGluT2-Cre ** p=0.0050 using ANOVA, mean ± SEM represented).

### NF-kB knockdown on VGat+ INs worsens immune dysfunction

To investigate the contribution of activation of Nf-kB after T9-SCI we manipulated IKKβ on VGat+ INs (IKKβ ^VGat^). Viral administration of AAV5-VGat-Cre resulted in more than half reduction in IKKβ in VGat+ INs (Fig. 5a, b). Mimicking the pattern of TNFR1 on VGat+ INs, IKKβ ^VGat^ was harmful to anti-viral immunity after chronic T9-SCI, leading to an increase in body weight loss (Fig. 5c). However, unlike TNFR1 manipulation, IKKβ ^VGat^ mice did not have significantly more viral load remaining in the lungs at 9 DPI or a significant decrease in spleen weight (Fig. 5d, e). However, IKKβ ^VGat^ immune dysfunction significantly reduces the amount of splenocytes and leukocytes (Fig. 5f, g) suggesting that NF-kB activity in these neurons is critical for proper neuro-immune communication to the spleen. However, TNFR1^VGat^ contributed to a more severe immune dysfunction after T9 SCI. Future studies will investigate alternative downstream pathways of TNFR1 (i.e., p38) and the additive effects of NF-κB and p38 knock-down on immune dysfunction after SCI.

**Figure 5.**
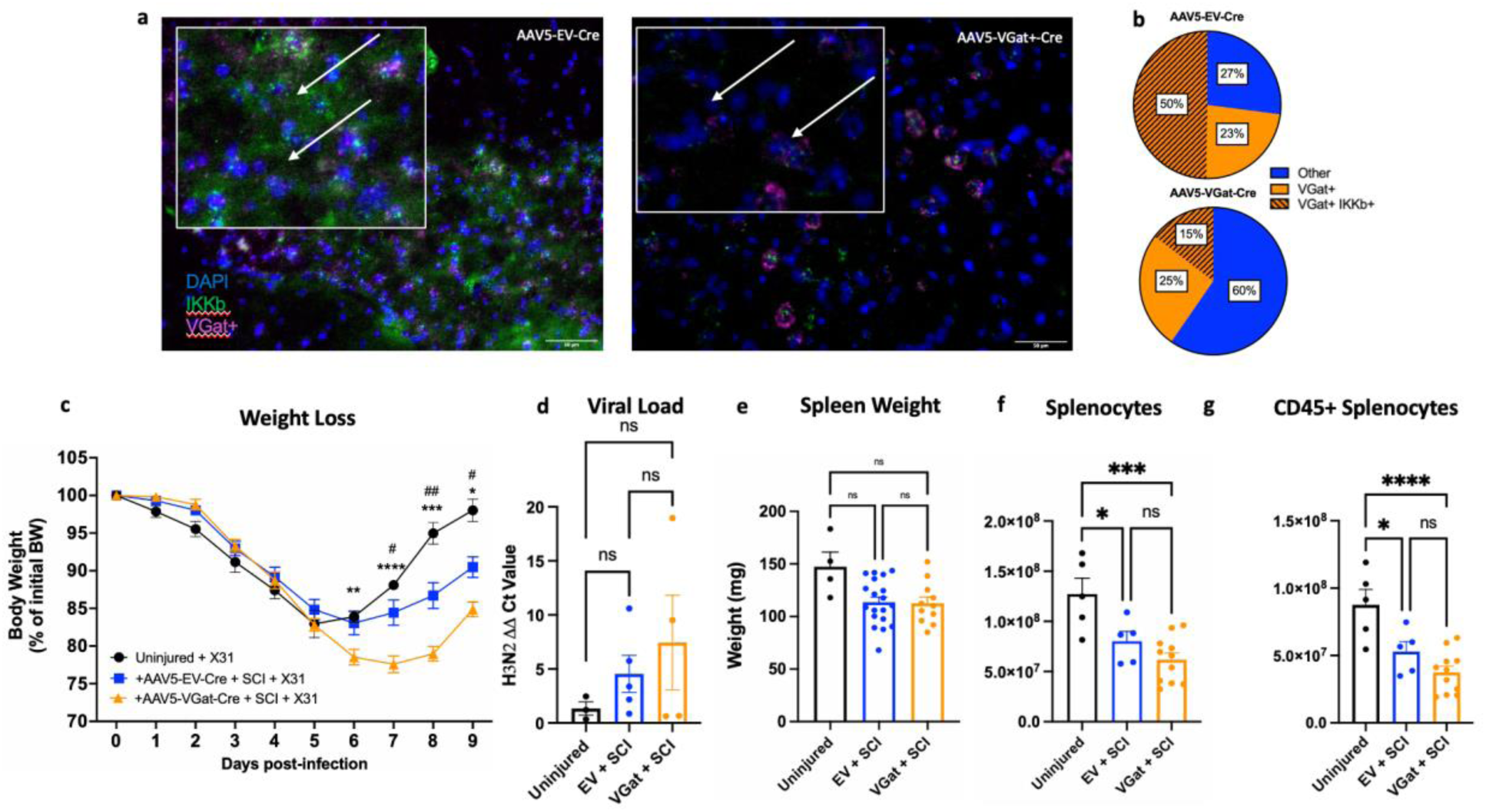
IKKβ KD on VGat+ INs does not improve anti-viral immunity after chronic T9-SCI. **a,b** Representative images of RNAscope analysis of VGat+ IKKβ + cells in the thoracic SC with AAV5-EV-Cre administration (b, top) or AAV5-VGat-Cre administration (b, bottom). Charts are made representing (orange) total VGat+ out of total (blue) and (dashes orange) VGat+ IKKβ +. **c** Body weight loss of mice for 9 DPI after chronic T9-SCI (* represent uninjured + X31 to AAV5-VGat-Cre, # represent AAV5-EV to VGat-Cre, Uninjured + X31 n=5 AAV5-EV-Cre n=20, AAV5-VGat-Cre n=12, 6 DPI ** p=0.0050, 7 DPI **** p= <0.0001 # p=0.016, 8 DPI *** p=0.0001, ## p=0.0029, 9 DPI *** p=0.0006 # p=0.0128 using ANOVA, mean ± SEM represented) **d** Viral load remaining in the lung at 9 DPI as determined by RT-qPCR for H3N2 viral load. **e, f** Spleen weight and total live splenocyte count of spleens from mice after 9 DPI. Weighed and analyzed at the time of sample collection and processing for flow cytometry analysis (splenocytes, uninjured to AAV5-EV-Cre * p=0.0282, AAV5-EV to VGat-Cre *** p=0.0003, using ANOVA, mean ± SEM represented) **g** Bar graph representing the total amount of CD45+ leukocytes in the spleen (uninjured to AAV5-EV-Cre * p=0.0211, AAV5-EV to VGat-Cre **** p=<0.0001).

## Discussion

Mitigating the effects of secondary conditions of individuals with SCI is crucial to decreasing mortality, and hospitalizations, and improving patients’ quality of life. Patients are repeatedly hospitalized and succumb to infections which continue to be the leading cause of death after SCI. Yet, there are currently no therapeutic strategies to mitigate the profound immune dysfunction after SCI. Our lab and others have reported that chronic T9-SCI negatively impacts the immune system and worsens anti-viral immunity. Unlike cervical and high-thoracic SCI, incomplete T9-SCI does not impact the innervation of the lungs via the pulmonary plexus or phrenic nerve, yet there is profound immune dysfunction in mice with T9-SCI when challenged with influenza, a moderate upper respiratory infection suggesting that organs innervated in the mid-thoracic spinal cord, i.e., spleen, are critical for anti-viral immunity.

Aside from the lung-draining lymph nodes which are crucial in priming effector CD8+ T cells, the spleen is capable of CD8+ T cell priming and virus specific CD8+ T cell activation independent of the lung-draining lymph node (Turner, Bickham, Farber, & Lefrançois, 2013). Additionally, in patients and animal models of SCI there are increased levels of proinflammatory cytokines which persist in the CNS weeks and months after injury. Chronic inflammation after injury and disease (i.e., SCI, TBI, Alzheimer’s, and MS) has previously been documented to alter neuron health and increase hyperexcitability of sympathetic circuitry in the CNS (Chen et al., 2024; Galvis-Montes et al., 2023; Ritzel et al., 2024). Moreover, targeting chronic inflammation after injury and disease has proven to be beneficial for disease progression and cognition (Dziedzic, Maciak, Miller, Starosta, & Saluk, 2024; Funaki, Nio-Kobayashi, Suzuki, & Bando, 2024). Therefore, we hypothesized that mitigating chronic inflammation in the CNS after SCI improves anti-viral immunity.

We report here that inhibiting pro-inflammatory sTNF in the spinal cord improves anti-viral immunity, increases virus-specific CD8 T cells in the lung, and decreases the residual viral load 9 days post-infection. Intrathecal administration of XPro1595 immediately or within the first few days of after SCI most beneficial for mitigating immune dysfunction due to the dramatic amounts of sTNF in the cord. In high-thoracic SCI, administration of XPro1595 after 2 weeks of SCI was not beneficial to mitigating secondary conditions (insert source: https://www.ncbi.nlm.nih.gov/pmc/articles/PMC8309421/). In addition, only intrathecal administration is neuroprotective and effective rather than systemic (insert source: https://www.ncbi.nlm.nih.gov/pmc/articles/PMC4176557/). Since a majority of the SCI field is focused on high-cervical and high-thoracic injuries, it is important to revisit these experiments in low-thoracic (T9) SCI and investigate the specific timeline of the effectiveness of XPro1595 after SCI to mitigate immune dysfunction.

Since XPro1595 was effective in improving anti-viral immunity when administered intrathecally we wanted to investigate how chronic inflammation on specific neuronal populations in the spinal cord could contribute to these effects. Our novel finding indicate that TNFR1 signaling on VGlut2^+^ excitatory interneurons contribute to immune dysfunction and supports previous work implicating spinal VGluT2 INs in disease (Testa et al., 2019; Zeng et al., 2020; Zhou & Roper, 2010; Ziemlińska et al., 2014) and secondary conditions of SCI (Noble et al., 2022; Ziemlińska et al., 2014). However, knock down of TNFR1 on VGat+ INs worsen anti-viral immunity supporting literature suggesting that inhibiting the excitability of inhibitory INs can cause an excitation/inhibition imbalance which may lead to profound excitability and neuronal damage.

It was intriguing that KD of TNFR1 on VGluT2+ INs improved antiviral immunity and spleen weight yet had minimal effects on total splenocytes and no differences on the abundance of T cells. Our lab previously reported that both T3 and T9-SCI increase levels of norepinephrine in the spleen which are decreased with intraspinal administration of XPro1595 suggesting that chronic inflammation in the spinal cord modulates the activity of the TH+ sympathetic nerve fibers innervating the spleen (Mironets et al., 2020). TH+ produced by nerve terminals in the spleen is the rate-limiting step in the synthesis of NE, and an increase in NE directly correlates to an increase in neuronal activity (Simkins et al., 2016). Thus, we hypothesize that chronic sTNF/TNFR1 activation on excitatory interneurons may increase the amount of NE in the spleen, contributing to more exhausted and/or less amounts of T cells in the spleen. Thereby, worsening response to infections after T9-SCI.

Although the mechanism is still unclear, our findings indicate that the effects we observed are NF-kB dependent but independent of the abundance of splenocytes and leukocytes. A limitation to this study is that all endpoint of immune cell analysis are 9 days post infection. Ongoing analysis aims to investigate how various steps of infection differ after T9-SCI. For example, altered antigen presentation, neutrophil activity and macrophage activity within the first few days of infection. Additional tools to investigate how NF-kB contributes to anti-viral immunity is by using a constitutively active IKKβ mouse model to investigate how persistent NF-kB activity will impact immune function with and without the presence of an injury. Our results indicate a that other inflammatory conditions which worsen immunity may benefit from XPro1595 or similar therapies. Alternatively, we cannot rule out that non-canonical pathways of TNFR1 activation such as p38 MAP Kinase also contribute to immune dysfunction after T9-SCI. p38 MAP Kinase activity has a reported role in immune response and chemotaxis so future studies will investigate if p38 activation increases with inflammation and if that is harmful to the sympathetic innervation of the spleen and immune dysfunction after T9-SCI. Moreover, immune cell exhaustion after chronic T9-SCI may contribute to worsened anti-viral immunity. Our lab has previously published that in correlation to an increase in NE there is an increase in exhausted immune cells in the spleen (Zha et al., 2014). *In vivo* NE stimulation proved to increase dysfunction of CD8+ T cells and an increase in PD-1. Future studies aim to investigate if TNFR1 activity on VGluT2+ and VGat+ affect the function of T cells and exhaustion in the spleen, including the effects of TH+ fiber outgrowth into the spleen and levels of NE.

Lastly, sTNF and TNFR1 hyper-activation has been reported in various diseases to contribute to pathogenesis and tissue damage (An et al., 2023; Ashenafi et al., 2023; Qi et al., 2023; Salai et al., 2023). Preliminary investigation if the effects of acute sTNF application on the intrinsic properties of vGluT2 and GAD67 spinal INs revealed minimal, varying, effects on cellular excitability (data not shown or supplementary figure ???). Both excitatory and inhibitory INs showed a significant increase in input resistance in the presence of sTNF. Although this can be indicative of an increase in cellular excitability, no other cellular parameters, including rheobase, significantly changed in the presence of sTNF. While we did not observe clear changes in the intrinsic properties of the spinal INs, the input to these neurons may be altered in the presence of sTNF (Park et al., 2011). It should also be noted that our recordings were performed in spinal cord slices from uninjured mice. It is possible that neuronal sensitivity to sTNF is altered post-SCI, as changes in the subtypes and expression levels of neuromodulatory receptors have been shown to lead to different effects on INs after SCI (Garcia-Ramirez, Ha, Bibu, Stachowski, & Dougherty, 2021; Husch et al., 2012). Studies are underway to better understand how SCI impacts the excitability of INs by sTNF.

Understanding how sTNF affects VGluT2+ interneurons will expand on novel therapeutic areas mitigating chronic inflammation after injury and other chronic inflammatory conditions. Ongoing work in our lab focuses on the effects of sTNF on VGluT2+ interneurons and its impact on nerve outgrowth into the spleen, contribution to NE and neuronal activity, immune cell function, and vasculature of the spleen, all of which may contribute to susceptibility to infection and mortality after T9-SCI.

## Conclusion

In summary, our results indicate that neuro-immune communication is altered after chronic T9-SCI and mitigating persistent activation of TNFR1 by sTNF using XPro1595 improves antiviral immunity. Specifically investigating TNFR1 on excitatory and inhibitory interneurons sheds light on the contribution of TNFR1 on excitatory interneurons in promoting immune dysfunction after T9-SCI. How TNF/TNFR1 in the cord impacts the spleen itself is still under investigation. Previous publications suggest that chronic T9-SCI increases the amount of norepinephrine in the spleen and the amount of exhausted immune cells. Future studies aim to investigate if TNFR1 knockdown on VGluT2+ prevents unnecessary immune cell exhaustion or decreases NE in the spleen, leading to an improvement in spleen health. Lastly, our report supports that the effects of persistent TNFR1 activation and worsened immune function occur dependent on NF-κB activation on both excitatory and inhibitory INs.

## Materials and Methods

### Mice

Mice were purchased from Jackson’s laboratory and bred in the animal facility at Drexel University. For these studies male and female C57Bl/6 wild-type (stock# 000664), IKKβ floxed (from M. Karin) and TNFR1 floxed (MGI, G. Kollias) were used. Electrophysiology experiments were performed using *vGlut2GFP* (kindly gifted by Dr. Ole Kiehn) (Caldeira, Dougherty, Borgius, & Kiehn, 2017)and *Gad67GFP* (Jax, #:003718; Oliva et al. 2000) transgenic mice. Mice were housed in groups of up to 5 mice with food and water *ad libitum* in a 12h light/dark cycle facility with controlled temperature and humidity. All experiments followed National Institutes of Health guidelines and were approved by Drexel University Institutional Animal Care and Use Committee.

### AAV5-Cre injections

Cell-specific recombination of floxed alleles specifically in the spinal cord was achieved by local delivery of the Cre recombinase using an AAV5 virus. AAV5-VGlut2-Cre and AAV5-VGat-Cre virus particles were obtained from Creative Biogene to specifically target VGlut2^+^ excitatory and VGat+ inhibitory interneurons respectively, promoters custom designed with Genecopoeia. AAV5 particles containing an empty vector were used as control. Adult mice (2 months old) were anesthetized using Ketamine (100 mg/Kg) and Xylazine (10 mg/Kg). Mice received a subcutaneous injection of Buprenorphine SR LAB (1 mg/Kg, ZooPharm) for analgesia. A laminectomy was performed at thoracic level T10-T12, and the animals were injected with the virus using a pre-pulled glass micropipette (World Precision Instrument) mounted onto a Hamilton syringe (1 ml of virus at manufacturer’s concentration of 1.00E+13 GC/ml (AAV5-VGluT2+-Cre) and 4.98E+9 GC/ml (AAV5-VGat-Cre) was slowly injected (0.4 ml/min) bi-laterally into the thoracic spinal cord.

### Spinal cord injury

Spinal cord injuries were performed as previously described (Bracchi-Ricard et al., 2016). Adult mice (3-4 months old) were deeply anesthetized with Ketamine (100 mg/Kg) and Xylazine (10 mg/Kg). Prior to surgery, animals were injected with Buprenorphine SR Lab (ZooPharm) for analgesia. A laminectomy was performed at thoracic level T9 to expose the spinal cord and injury was induced using the Infinite Horizon impactor (Precision Systems) set at a predetermined force (65 kDynes) which generates a displacement of the spinal cord around 500um. After suturing, the mice received 1 mL lactated ringer solution for hydration and gentamicin (5 mg/Kg) Bladders were expressed twice daily, and gentamicin given daily for a week to prevent urinary tract infections.

### Influenza virus infection

Influenza virus subtype A (HKx31, H3N2) was purchased from Charles River (material number 10100375). Intranasal delivery of the virus was performed as previously described (Bracchi-Ricard et al., 2016). In brief, mice were anesthetized with Ketamine (100 mg/Kg) and Xylazine (10 mg/Kg), placed on their back and 20 ml of virus suspension (25 HAU) were applied in the nostrils.

### Drug treatment

The anti-TNF biologics, Xpro1595 (INmune Bio) was administered directly to the site of injury using Alzet osmotic pump (model 1004) with Brain infusion Kit 3 (Alzet). Rate of delivery and amount of XPro1595 loaded in the pump was 2.5 mg/mL concentration/1 μL/h.

### Viral load assessment

To assess the levels of influenza virus we performed quantitative PCR for the viral matrix protein M1 gene on the infected lungs (Bracchi-Ricard et al., 2016). In brief, total RNA was isolated from the lungs of influenza infected mice using Trizol™ Reagent (InVitrogen). cDNAs were obtained by reverse transcription using Omniscript RT kit (Qiagen).

Quantitative real-time PCR was done with the QuantiFast SYBR Green kit (Qiagen) on the RotorGene (Qiagen) with the following primers: M1 (virus-specific matrix protein M1 gene) *forward* 5’-AGCCGAGATCGCGCAGAGACT, M1 *reverse* 5’-TGAGCGTGAACACAAATCCTAAAA-3’, b-actin *forward* 5’-ATGGTGGGAATGGGTCAGA -3’, b-actin *reverse* 5’-CACGCAGCTCATTGTAGAAGG-3’. Quantification was done utilizing the delta-delta Ct method to measure the relative change in gene expression.

### Cell isolation from spleen and lung

Mice were euthanized with CO_2_ and their spleen and lungs quickly removed and stored on ice in RPMI1640 medium (Quality Biologicals). Spleens were dissociated to single cell suspension through a 40-mm filter. Red blood cells (RBCs) were lyzed using RBC lysis buffer (Biolegend) for 3 min at room temperature. Lung tissues were cut into small pieces and then digested in RPMI1640 containing 3 mg/mL of collagenase A (Roche) and 0.15 mg/mL of DNase I (Roche) for 90 min in a 37°C/5%CO_2_ incubator. Next, the tissue was passed through a 70-mm filter. Red blood cell lysis was performed as described above. Finally, both spleen and lung cell pellets were resuspended in MACS buffer (0.5% BSA.2mM EDTA in PBS) to a cell concentration of 2×10^7^ cells/ml.

### Flow cytometry

Cells were stained with the Live/Dead™ Fixable Aqua stain according to manufacturer’s protocol (ThermoFisher). Then aliquots of 100 ml (2×10^6^) cells were plated in a 96-well round-bottom plate (Corning, catalog 3797). Cells were stained with the following antibody mixes for 1 hour at 4°C in the dark: Mix 1: AlexaFluor 700 anti-CD45 (1:150, BD Biosciences), AlexaFluor 647 anti-CD44 (1:100, Biolegend), H-2Db Influenza NP Tetramer-ASNENMETM-PE (1:100, MBL International), AlexaFluor 488 anti-CD8a (1:100, Biolegend), BV785 anti-CD4 (1:400, Biolegend), BV650 anti-CD62L (1:200, BD Biosciences). Mix 2: AlexaFluor 700 anti-CD45 (1:150, BD Biosciences), AlexaFluor 647 anti-CD44 (1:100, Biolegend), H-2Db Influenza PA Tetramer-SSLENFRAYV-PE (1:100, MBL International), AlexaFluor 488 anti-CD8a (1:100, Biolegend), BV785 anti-CD4 (1:400, Biolegend), BV650 anti-CD62L (1:200, BD Biosciences). Mix 3: AlexaFluor 700 anti-CD45 (1:150, BD Biosciences), PE anti-CD11b (1:150, Biolegend), APC anti-Ly6G (1:75, Biolegend), FITC anti-CD45R/B220 (1:100, BD Biosciences), BV605 anti-CD161 (1:100, Biolegend). Mix 4: AlexaFluor 700 anti-CD45 (1:150, BD Biosciences), BB515 anti-CD11b (1:100, BD Biosciences), BV650 anti-CD11c (1:100, Biolegend), PE anti-MHCII (I-A, 1:50, ThermoFisher), PE-Cy7 anti-Ly6c (1:75, Biolegend). Samples were analyzed on a BD FACSymphony™ A3 or A5 SE II analyzer (BD Biosciences) at the Wistar Institute Flow Cytometry Facility (Philadelphia, PA). All data was analyzed using FlowJo^TM^ v10.8 Software (BD Lifesciences). First, all debris and doublets were excluded. Then, live and CD45+ cells were selected from gating strategy. Cells were further gated based on cell type of interest. CD4+ T cells (CD45+, CD8-CD4+), CD8+ T cells (CD45+CD4-CD8+), activated T cells (CD45+CD8+CD44+CD62+), NP/PA+ virus specific CD8+ T cells (CD45+ CD4-CD8+NP+/PA+).

### RNAscope

Mice were deeply anesthetized with ketamine and xylazine prior to perfusing with cold PBS to remove blood. The mice were then perfused with 10% Neutral Buffered Formalin (NBF) and spinal cords were harvested and stored in 10% NBF overnight. Following cryoprotection by incubation in a series of 40% then 30% sucrose/PBS solution overnight at 4°C, the spinal cords were then embedded in Tissue-Tek® O.C.T. compound (Sakura) and cut on a cryostat. Serial sections (15 µm thickness, 7 series) were collected on positively charged microscope slides (VWR) and stored at –80°C until further processing. RNAscope processing followed ACDBio RNAScope ® Multiplex Fluorescent Assay manufacturer’s guidelines using probes Mm-TNFRsf1a (cat# 438941-C1) or Mm-Ikbkb (cat# 549481-C1), MmSlc17a6 (cat# 456751-C2), and Mm-SLc32a1 (cat# 319191-C3) from ACDBio. Images were taken using the Zeiss AxioObserver at the CIC Core at Drexel University Department of Biology and analysis was done using Image J. Quantification was accomplished by counting total number of cells (DAPI) followed by co-localization with TNFR1, IκκB, VGluT2, and VGat.

## Acknowledgements

We thank Mr. Jeffrey Faust for expert technical and experimental guidance with flow cytometry and Dr. Harini Sreenivasappa for expert assistance in confocal imaging at Drexel’s Cell Imaging Center.

